# Evaluating the performance of 3-tissue constrained spherical deconvolution pipelines for within-tumor tractography

**DOI:** 10.1101/629873

**Authors:** Hannelore Aerts, Thijs Dhollander, Daniele Marinazzo

## Abstract

The use of diffusion MRI (dMRI) for assisting in the planning of neurosurgery has become increasingly common practice, allowing to non-invasively map white matter pathways via tractography techniques. Limitations of earlier pipelines based on the diffusion tensor imaging (DTI) model have since been revealed and improvements were made possible by constrained spherical deconvolution (CSD) pipelines. CSD allows to resolve a full white matter (WM) fiber orientation distribution (FOD), which can describe so-called “crossing fibers”: complex local geometries of WM tracts, which DTI fails to model. This was found to have a profound impact on tractography results, with substantial implications for presurgical decision making and planning. More recently, CSD itself has been extended to allow for modeling of other tissue compartments in addition to the WM FOD, typically resulting in a 3-tissue CSD model. It seems likely this may improve the capability to resolve WM FODs in the presence of infiltrating tumor tissue. In this work, we evaluated the performance of 3-tissue CSD pipelines, with a focus on within-tumor tractography. We found that a technique named single-shell 3-tissue CSD (SS3T-CSD) successfully allowed tractography within infiltrating gliomas, without increasing existing single-shell dMRI acquisition requirements.

## Introduction

The goal of neurosurgery for brain tumors is to maximally remove harmful tumor tissue, while minimizing the risk of inducing permanent neurological deficits which might be caused by damaging healthy functioning tissue. This pertains especially to glioma tumors, as they often grow by infiltration of such healthy tissue. As a result, white matter tracts can effectively be present *within* a tumor or in immediately adjacent brain tissue (Skirboll et al., 1996), that might also show a degree of partial voluming with the tumor region. In addition to infiltration, white matter pathways can also be displaced due to so-called mass effects, or disrupted in the case of complete infiltration (Campanella et al., 2014; Essayed et al., 2017). Therefore, it is of utmost importance to identify the location and extent of white matter tracts in the tumor area that should be maximally preserved during surgery, in a patient-specific manner.

To this end, diffusion MRI (dMRI) guided fiber tractography is often employed in presurgical planning (Dimou et al., 2013). dMRI is the only modality currently available that allows non-invasive assessment of tissue microstructure in-vivo, and without the use of ionizing radiation. In particular, dMRI measurements allow for the estimation and modeling of the local orientations of white matter tracts. Subsequently, tractography algorithms trace this information throughout the brain to infer long-range connectivity between distant brain regions.

Originally, in the majority of neurosurgical dMRI studies as well as in clinical practice, the diffusion tensor imaging (DTI) model (Basser et al., 1994) was typically used to estimate the white matter fiber orientation in each voxel from the dMRI data (Essayed et al., 2017; Farquharson et al., 2013; Nimsky et al., 2016). DTI-based tractography methods are however fundamentally (and *a priori*) limited because the DTI model can only resolve a single fiber direction within each imaging voxel, whereas it has been shown that up to 90% of white matter voxels in the brain contain more complex architectures, typically referred to as “*crossing fibers*” (Jeurissen et al., 2013). In such regions, the single orientation obtained from DTI is unreliable and may cause both anatomically implausible (false positive) as well as missing (false negative) tracts (Farquharson et al., 2013; Jeurissen et al., 2013). Both types of bias can have detrimental consequences in the context of presurgical planning (Duffau, 2014a; Duffau, 2014b; Nimsky et al., 2016). Overestimation of white matter tracts can lead to incomplete resection of the tumor, which can significantly diminish patients’ survival rates after neurosurgery (Kramm et al., 2006). Underestimation of white matter tracts close to the lesion, on the other hand, can lead to erroneous removal of parts of healthy tracts, which can cause functional impairments. To address the *crossing fiber* problem, several higher-order models have been developed; see (Tournier et al., 2011) for a review. One such frequently adopted higher-order approach is constrained spherical deconvolution (CSD) (Tournier et al., 2007). In short, CSD uses high angular resolution diffusion imaging (HARDI) (Tuch et al., 2002) data to generate estimates of a full continuous angular distribution of white matter (WM) fiber orientations within each imaging voxel, without requiring prior knowledge regarding the number of fibers in any given voxel (Tournier et al., 2007). Generally, such higher-order approaches have been shown to yield superior results compared to the original tensor model (Neher et al., 2015). In addition, application of CSD in presurgical planning has been demonstrated to result in superior determination of location and extent of WM tracts compared to the DTI model, hence providing improved estimates of safety margins for neurosurgical procedures (Farquharson et al., 2013; Küpper et al., 2015; Mormina et al., 2016).

Recently, a multi-tissue extension of CSD has been proposed, that allows accurate modelling of not only pure white matter (WM) voxels, but also voxels (partially) containing gray matter (GM) and cerebrospinal fluid (CSF) (Jeurissen et al., 2014). By including such additional compartments in the model to explicitly account for signal arising from non-WM tissue^1^ types, multi-shell multi-tissue CSD (MSMT-CSD) allows for more accurate estimation of the WM fiber orientation distributions (FODs) as well: the signal originating from other tissue types otherwise contaminates these WM FODs when using the original “single-tissue” CSD (ST-CSD), resulting in severely distorted WM FOD shapes. The most important drawback of MSMT-CSD, compared to the original ST-CSD, is that it requires dMRI data acquired with multiple b-values, commonly referred to as “multi-shell” dMRI data. Specifically, for MSMT-CSD to resolve the typical triplet of tissues (WM, GM and CSF), it requires at least 3 *different* b-values (including non-diffusion weighted b=0 data). Advanced acquisition protocols to obtain such data are however not always readily available in clinical MRI facilities, and even if they would be, acquisition of multi-shell data is more time consuming, limiting their practical usefulness. Moreover, it also poses greater challenges for several preprocessing steps, such as motion and distortion correction, as each unique b-value results in a different contrast.

In an attempt to overcome these practical limitations and avoid the advanced acquisition requirements associated with MSMT-CSD, a method called single-shell 3-tissue CSD (SS3T-CSD) was recently proposed (Dhollander and Connelly, 2016; Dhollander et al., 2016). This method aims to obtain similar results compared to MSMT-CSD, yet by using only *single-shell* (+b=0) data to model the same tissue compartments (WM, GM and CSF). It has also been shown to be able to fit other, e.g. pathological, tissue compositions by using the same set of WM, GM and CSF tissue compartments (Dhollander et al., 2017). In the context of CSD techniques, the basic model for each tissue compartment—also referred to as its response function—is typically estimated from the data themselves beforehand by identifying voxels containing more “pure” samples of these respective tissue types. To avoid confusion, when using the typical triplet of response functions derived from *actual* WM, GM and CSF voxels to model *any* (potentially pathological) other tissue composition, a terminology of so-called “WM-like”, “GM-like” and “CSF-like” tissues was introduced (Dhollander et al., 2017). It was for instance shown that white matter hyperintense lesions in Alzheimer’s disease patients contain an amount of “GM-like” tissue. This does of course not mean that these lesions are biologically similar to genuine GM in any way, but rather that (part of) the dMRI signal measured in the lesion resembles the dMRI signal also observed in GM. This principle was subsequently successfully leveraged to study heterogeneity within lesions in Alzheimer’s disease and differentiate (parts of) such lesions according to their WM-, GM- and CSF-like composition (Mito et al., 2018; Mito et al., 2019).

An overview of the aforementioned CSD techniques is provided in Table 1. Note that the term “ST-CSD”, as introduced above, is merely used as a synonym for the “original” CSD technique (Tournier et al., 2007), yet communicates more explicitly what kind of CSD technique (i.e., *single-tissue*) it is referring to.

**Table 1.** Overview of CSD techniques.

|  | **“Single-Tissue” CSD**<br>**(ST-CSD)**<br>(Tournier et al., 2007) | **Multi-Shell Multi-Tissue CSD**<br>**(MSMT-CSD)**<br>(Jeurissen et al., 2014) | **Single-Shell 3-Tissue CSD**<br>**(SS3T-CSD)**<br>(Dhollander & Connelly, 2016;<br>Dhollander et al., 2016) |
| --- | --- | --- | --- |
| **Acquisition requirements** | Single shell<br>Preferably high angular resolution and a high b-value | Multiple shells<br>At least 3 unique b-values (including b=0) to account for 3 tissue types | Single shell (+b=0)<br>b=0 is included in *any* typical (single-shell) dMRI acquisition |
| **Modeling capabilities** | “Pure” normal WM only<br>In case of partial voluming with other tissue types, the WM FOD is distorted by their signal | WM (FOD) and GM/CSF compartments<br>These might account for other (pathological) tissue compositions too | WM (FOD) and GM/CSF compartments<br>These might account for other (pathological) tissue compositions too |

In this work, we investigated the possible benefits of 3-tissue CSD techniques (MSMT-CSD and SS3T-CSD) over the original single-tissue CSD in reconstructing white matter tracts close to, but especially within, infiltrative brain tumors. In addition, we evaluated the relative performance of MSMT-CSD and SS3T-CSD for this purpose, to determine whether the greater acquisition requirements of MSMT-CSD can be justified by the resulting white matter tract reconstruction. To this end, multi-shell dMRI data were acquired from seven glioma patients on the day before tumor resection. WM fiber orientation distributions (FODs) were computed for all patients using a ST-CSD, MSMT-CSD and SS3T-CSD pipeline. We focused on qualitative comparison of the WM FODs within (and close to) the tumor region. Finally, we performed tractography on all outcomes to directly assess the impact of the different CSD techniques on the quality of tract reconstruction.

## Methods

### Participants

In this study we included patients who were diagnosed with a glioma; a primary brain tumor developing from glial cells (Fisher et al., 2007). Gliomas are typically classified based on their malignancy using the World Health Organization grading system, where grade I tumors are least malignant and grade IV tumors are most malignant. Hereby, “malignancy” relates to multiple aspects: the speed at which the disease evolves, the extent to which the tumor infiltrates healthy brain tissue, and chances of recurrence or progression to higher grades of malignancy.

Patients were recruited from Ghent University Hospital (Belgium) between May 2015 and October 2017. Patients were eligible if they (1) were at least 18 years old, (2) had a supratentorial glioma WHO grade II or III for which a surgical resection was planned, and (3) were medically approved to undergo MRI investigation. Seven patients meeting these inclusion criteria (mean age 50.7y, standard deviation = 11.7; 43% females; patient characteristics are described in Table 2) were identified. Testing took place at the Ghent University Hospital on the day before each patient’s surgery. All participants received detailed study information and gave written informed consent prior to study enrollment. This research was approved by the Ethics Committee of Ghent University Hospital (approval number B670201318740). All neuroimaging data used for this study are publicly available at the OpenNeuro website (https://openneuro.org/) and on the European Network for Brain Imaging of Tumours (ENBIT) repository (https://www.enbit.ac.uk/) under the name “BTC_preop”.

**Table 2.** Patient characteristics.

| Subject ID | Sex | Age | Tumor lateralization | Tumor location | Tumor size (cm<sup>3</sup>) | Tumor histology* |
| --- | --- | --- | --- | --- | --- | --- |
| PAT05 | F | 40 | Left | Frontal | 12.95 | Oligo-astrocytoma II |
| PAT07 | M | 55 | Left | Temporal | 33.94 | Ependymoma II |
| PAT16 | M | 39 | Right | Fronto-temporal | 50.24 | Anaplastic astrocytoma II-III |
| PAT20 | F | 70 | Right | Parietal | 15.24 | Anaplastic astrocytoma III |
| PAT25 | F | 46 | Right | Temporal | 18.56 | Glioma II |
| PAT26 | M | 61 | Right | Temporal | 59.21 | Anaplastic astrocytoma III |
| PAT28 | M | 44 | Left | Frontal | 11.49 | Oligodendroglioma II |

\* (*Oligo-*)*astrocytoma*, *ependymoma*, and *oligodendroglioma* are subtypes of glioma tumors.

### MRI data acquisition

From all participants, MRI scans were obtained using a Siemens 3T Magnetom Trio MRI scanner with a 32-channel head coil. First, a T1-weighted MPRAGE anatomical image was acquired: 160 slices, TR = 1750 ms, TE = 4.18 ms, field of view = 256 mm, flip angle = 9°, voxel size 1 x 1 x 1 mm. The total acquisition time was 4:05 min. Further, a multi-shell high-angular resolution diffusion imaging (HARDI) scan was acquired with the following parameters: a voxel size of 2.5 x 2.5 x 2.5 mm, 60 slices, TR = 8700 ms, TE = 110 ms, field of view = 240 mm. The dMRI acquisition consisted of 102 volumes in total: 6 non-diffusion weighted (b=0) volumes, and respectively 16, 30 and 50 gradient directions for b = 700, 1200 and 2800 s/mm^2^. The total acquisition time for the complete multi-shell protocol was 15:14 min. In addition, two separate non-diffusion weighted (b=0) images were acquired with reversed phase-encoding blips to correct for susceptibility induced distortions (Andersson et al., 2003).

### Diffusion MRI processing

#### Preprocessing

Diffusion MRI data were preprocessed using a combination of FSL (FMRIB Software Library) (Jenkinson et al., 2012) and MRtrix3 (Tournier et al., 2019). Specifically, the dMRI data were denoised (Veraart et al., 2016), corrected for Gibbs ringing artifacts (Kellner et al., 2016), motion and eddy currents (Andersson and Sotiropoulos, 2016), susceptibility induced distortions (Andersson et al., 2003), and bias field induced intensity inhomogeneities (Zhang et al., 2001). Brain masks were computed using the Brain Extraction Tool (Smith, 2002). Next, each subject’s high-resolution anatomical image was linearly registered to the dMRI data using FSL FLIRT (Jenkinson et al., 2002; Jenkinson and Smith, 2001).

#### Response function estimation and CSD modeling pipelines

##### ST-CSD

Only the highest b-value (b = 2800 s/mm^2^) shell was used for ST-CSD processing, in line with typical recommendations (Tournier et al., 2013): the high b-value data typically has better contrast-to-noise properties which, together with a high angular resolution, are optimally suited for the requirements of ST-CSD. The single-fiber WM response function was estimated from the data themselves using an iterative approach (Tournier et al., 2013). Finally, using this WM response function, ST-CSD was performed to obtain the WM FODs in all voxels (Tournier et al., 2007).

##### MSMT-CSD

The complete multi-shell dataset (all b-values) was used for MSMT-CSD processing: this gradient scheme (i.e., b-values and numbers of gradient directions per individual b-value) was very similar to the one used in (Jeurissen et al., 2014), with increasing numbers of gradient directions for higher b-values. The multi-shell single-fiber WM response function as well as multi-shell isotropic GM and CSF response functions were estimated from the data themselves using a T1-weighted image segmentation guided method (Jeurissen et al., 2014). Finally, using these 3 (WM, GM, CSF) response functions, MSMT-CSD was performed to obtain the WM FODs as well as GM and CSF compartments in all voxels (Jeurissen et al., 2014).

##### SS3T-CSD

Only the highest b-value (b = 2800 s/mm^2^) shell and the b=0 data were used for SS3T-CSD processing (Dhollander and Connelly, 2016), which ensures optimal contrast-to-noise properties within the shell (similar to ST-CSD recommendations) as well as between the single-shell and the b=0 data. The single-shell (+b=0) single-fiber WM response function as well as single-shell (+b=0) isotropic GM and CSF response functions were estimated from the data themselves using a fully automated unsupervised method (Dhollander et al., 2016; Dhollander et al., 2019). Finally, using these 3 (WM, GM, CSF) response functions, SS3T-CSD was performed to obtain the WM FODs as well as GM and CSF compartments in all voxels (Dhollander and Connelly, 2016).

#### Tractography

Probabilistic whole-brain fiber tractography was performed for each subject and guided by the WM FOD image resulting from each of the three different CSD techniques (ST-CSD, MSMT-CSD, SS3T-CSD) using the “second order integration over fiber orientation distributions” (iFOD2) algorithm (Tournier et al., 2010). Tractography was seeded across the entire brain volume and 500,000 streamlines were generated using an FOD amplitude threshold of 0.07 as a stopping criterion, to avoid tracking small noisy features of FODs which might be unrelated to genuine WM structure.

For both 3-tissue CSD techniques (MSMT-CSD and SS3T-CSD), we expected WM FODs to be smaller in the tumor regions, reflecting a reduced presence of healthy axons due to infiltration of tumor tissue (as diffusion signal resulting from the tumor tissue might be “picked up” by the non-WM compartments in the model instead) and other potential sources of WM damage. To address this challenge, we devised a pragmatic solution where we gradually reduced the FOD amplitude threshold close to and even more so within the tumor. To this end, we first registered the T1-weighted image to the dMRI data using FSL’s registration tools (FLIRT) (Jenkinson et al., 2002; Jenkinson and Smith, 2001). Next, tumors were manually delineated based on the T1-weighted images, and further automatically optimized using the Unified Segmentation with Lesion toolbox (Phillips and Pernet, 2017). These tumor segmentations were then spatially smoothed using a Gaussian kernel with a standard deviation of 3 mm, to introduce a smooth boundary extending slightly beyond—as well as within—the edges of the tumor. Finally, during the actual tractography process, the FOD amplitude threshold was reduced by up to a factor 3 within the tumor, modulated by the smoothed tumor segmentation.

## Results

### Tissues and FODs from different CSD modelling pipelines

CSD results for four representative patients (PAT05; PAT16; PAT26; PAT28) are shown in Figures 1–4. In each of these figures, the columns provide a direct comparison between the outcomes of the three different CSD modelling pipelines defined earlier in the Methods section.

**Figure 1.**
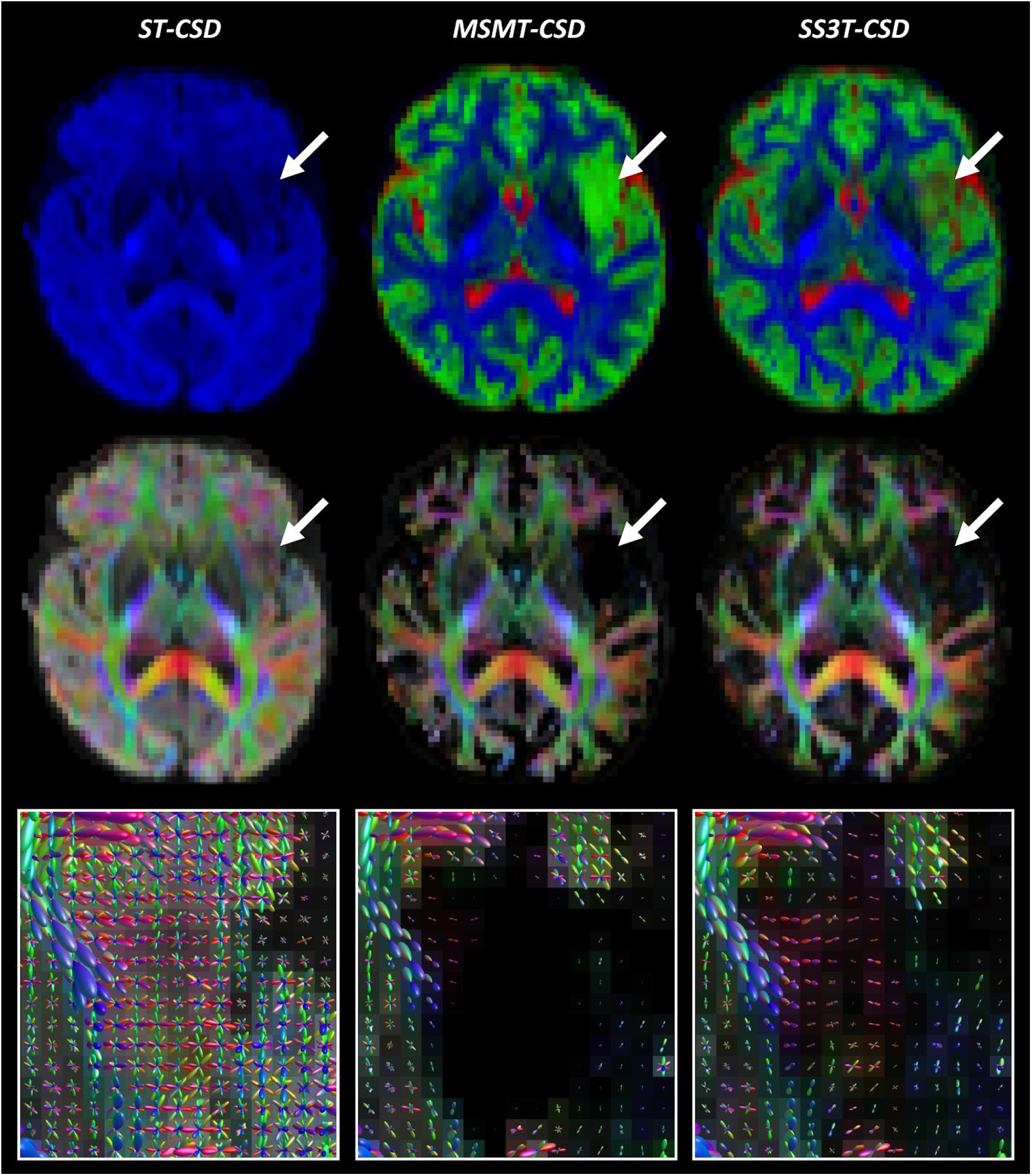
Comparison of ST-CSD, MSMT-CSD and SS3T-CSD outcomes for patient PAT05 (oligo-astrocytoma WHO grade II). Top row: tissue-encoded color maps (*blue*: WM-like; *green*: GM-like; *red*: CSF-like). Middle row: WM FOD-based directionally-encoded color (DEC) maps (*red*: left-right; *green*: anterior-posterior; *blue*: superior-inferior). Bottom row: WM FODs overlaid on FOD-based DEC map within tumor region.

**Figure 2.**
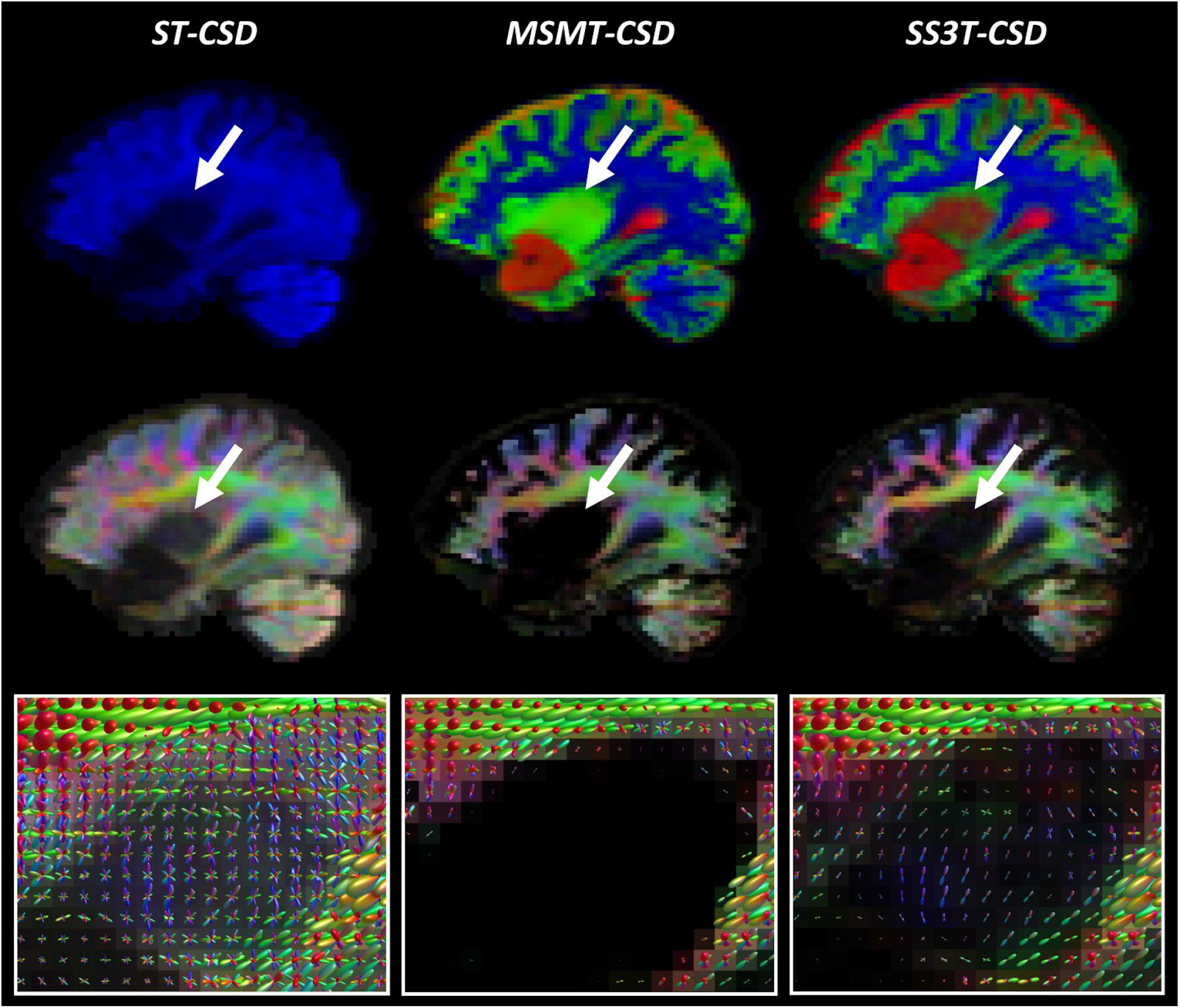
Comparison of ST-CSD, MSMT-CSD and SS3T-CSD outcomes for patient PAT16 (anaplastic astrocytoma WHO grade II-III). Top row: tissue-encoded color maps (*blue*: WM-like; *green*: GM-like; *red*: CSF-like). Middle row: WM FOD-based directionally-encoded color (DEC) maps (*red*: left-right; *green*: anterior-posterior; *blue*: superior-inferior). Bottom row: WM FODs overlaid on FOD-based DEC map within tumor region.

**Figure 3.**
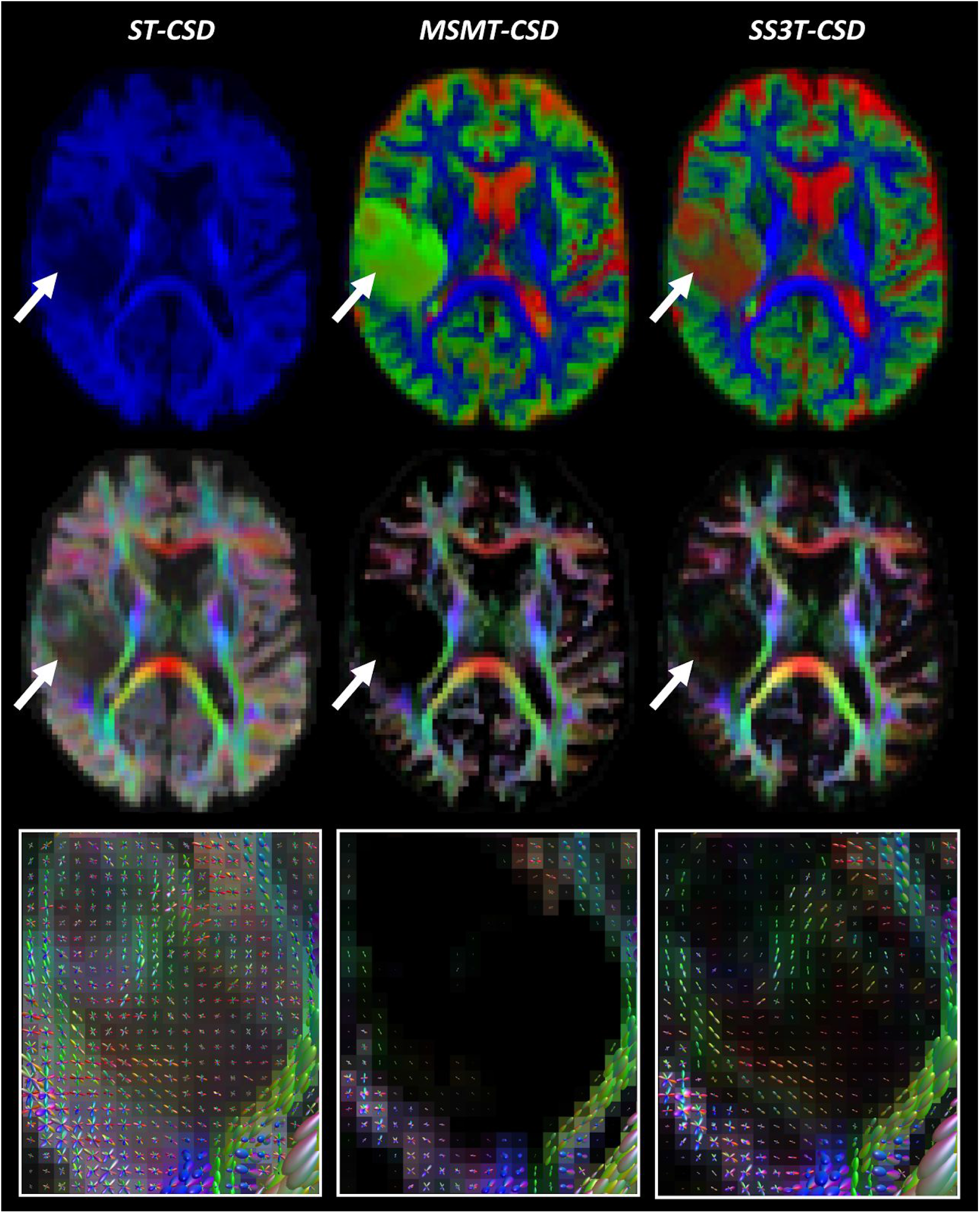
Comparison of ST-CSD, MSMT-CSD and SS3T-CSD outcomes for patient PAT26 (anaplastic astrocytoma WHO grade III). Top row: tissue-encoded color maps (*blue*: WM-like; green: GM-like; *red*: CSF-like). Middle row: WM FOD-based directionally-encoded color (DEC) maps (*red*: left-right; *green*: anterior-posterior; *blue*: superior-inferior). Bottom row: WM FODs overlaid on FOD-based DEC map within tumor region.

**Figure 4.**
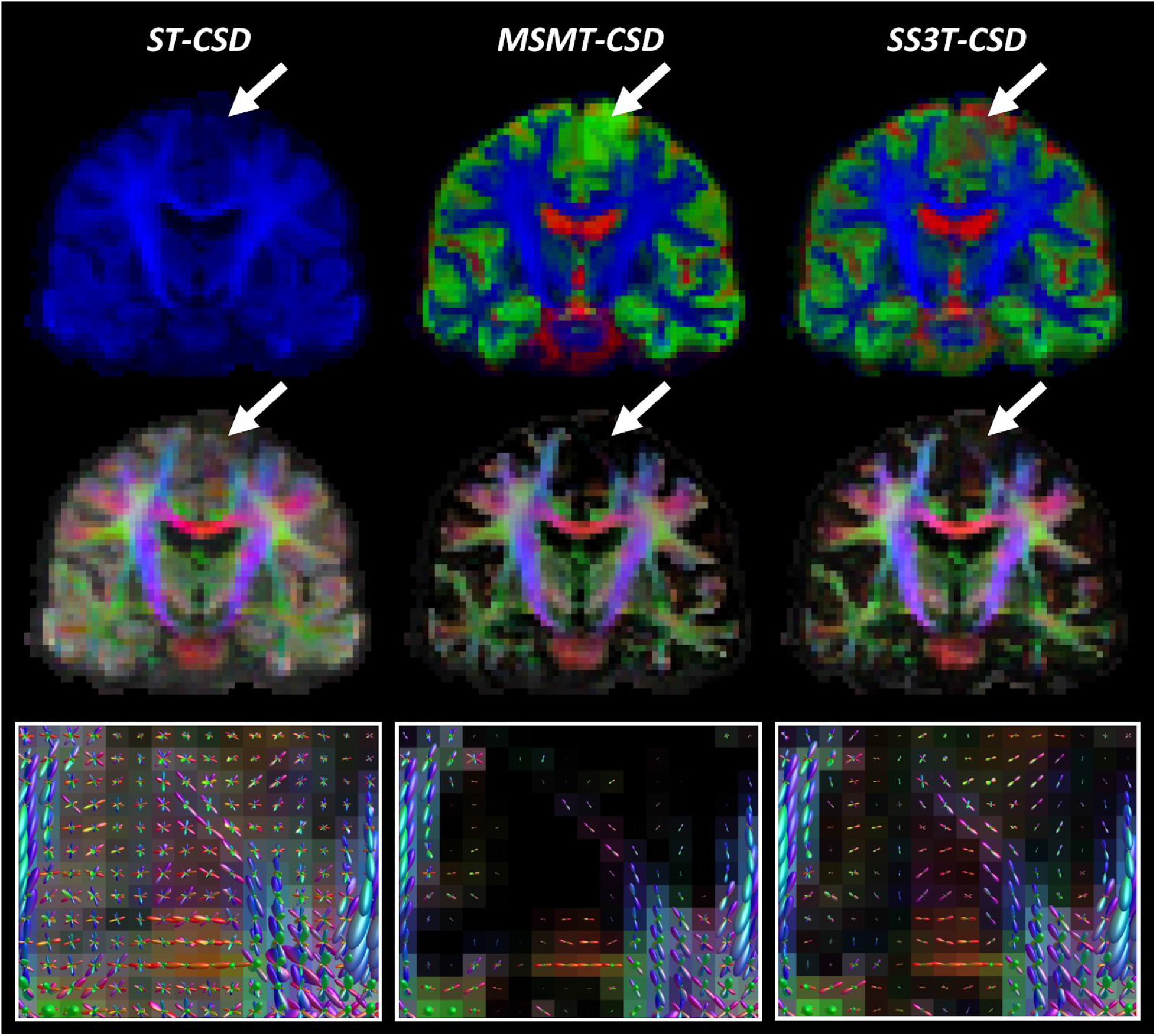
Comparison of ST-CSD, MSMT-CSD and SS3T-CSD outcomes for patient PAT28 (oligodendroglioma WHO grade II). Top row: tissue-encoded color maps (*blue*: WM-like; *green*: GM-like; *red*: CSF-like). Middle row: WM FOD-based directionally-encoded color (DEC) maps (*red*: left-right; *green*: anterior-posterior; *blue*: superior-inferior). Bottom row: WM FODs overlaid on FOD-based DEC map within tumor region.

The top rows of Figures 1–4 depict tissue encoded color images (Dhollander et al., 2018; Jeurissen et al., 2014), where *blue* encodes the WM-like compartment, *green* the GM-like compartment and *red* the CSF-like compartment. The tumor region in each of these figures is indicated by an arrow. Naturally, ST-CSD results only include a WM-like (*blue*) compartment, which accumulates all dMRI signal (of the b = 2800s/mm^2^ shell) regardless of which combination of tissues it might have originated from. In contrast, MSMT-CSD and SS3T-CSD results both feature all 3 tissue compartments, but show a consistent difference between both methods that stands out particularly clearly in regions where tumor tissue is present. That is, in MSMT-CSD results, the GM-like (*green*) compartment appears to dominate strongly across the entire tumor, whereas in SS3T-CSD results it is generally less strong, in favor of CSF-like and supposedly WM-like contributions (although the latter are hard to visually observe on the tissue maps). In addition, SS3T-CSD results show spatially varying tissue compositions throughout the tumors, with the strongest GM-like contributions still appearing in genuine cortical GM nearby the tumor or infiltrated by it. Finally, note that the result of PAT16 also shows a large fluid filled cavity directly below the tumor, resulting from a prior resection.

The middle row in each figure shows FOD-based directionally-encoded color (DEC) maps (Dhollander et al., 2015), where colors encode the local orientations of WM-like structures using a common convention used to display dMRI results (*red*: left-right; *green*: anterior-posterior; *blue*: superior-inferior). Note that these FOD-based DEC maps are different from traditional DTI-based DEC fractional anisotropy (FA) maps. While the color of DEC FA maps is determined only by the single main tensor orientation, FOD-based DEC maps derive their color from all orientations of the full FOD. Also, the intensity of the FOD-based DEC map is directly proportional to the WM-like compartment itself (i.e., equivalent to the *blue* color in the tissue encoded color images in the first row). Again, the most obvious difference is observed between ST-CSD on the one hand and both 3-tissue CSD techniques on the other hand: due to ST-CSD capturing all signal in its WM-like compartment, intensities are high throughout the whole brain parenchyma, including the cortical GM. The (WM-like) intensity is reduced to varying degrees in several tumor regions though. In both MSMT-CSD and SS3T-CSD results, GM-like and CSF-like signals are filtered out, leaving only genuine WM. This is also strongly the case in all tumors, which appear to be very low in (healthy) WM-like content. In the MSMT-CSD results, WM-like signal was effectively entirely absent (i.e., strictly zero) in many tumor voxels. While hard to visually observe on the FOD-based DEC maps, in SS3T-CSD, traces of WM-like signal remain for most tumor voxels. The actual WM FODs which represent these WM-like signals provide more insight.

Finally, the bottom rows of Figures 1–4 show the WM FODs themselves, directly overlaid on their corresponding FOD-based DEC map, for a region zoomed in on the tumor’s location (i.e., the region indicated by the arrow in the other rows). For the purpose of clear visualization and optimal assessment of the FODs, all FODs are scaled by the same factor across all results, so as to remain consistent and directly comparable between results. Looking at the FODs in the tumors, differences between all three CSD pipelines are most apparent. ST-CSD results show WM FODs in the tumor areas, but these are very noisy and show little to no spatial coherence. This is similar to the WM FODs from ST-CSD found in cortical GM, which are known to feature many false positive noisy lobes as well, due to presence of non-WM tissue (Jeurissen et al., 2014). MSMT-CSD results, remarkably, demonstrate the opposite effect: as most to all WM-like signal is absent in many tumor voxels, these voxels show almost or entirely no WM FODs. Perhaps surprisingly, SS3T-CSD does show WM FODs in most of these voxels. Moreover, the FODs from SS3T-CSD reveal a spatially coherent pattern, which also appears to “connect” well to nearby WM FODs in healthy white matter outside of the tumors. Due to the strongly reduced amount of WM-like signal though, SS3T-CSD FODs inside the tumor are typically reduced substantially in amplitude. The latter observation motivated our choice to introduce a mechanism which reduces the FOD amplitude threshold used as a stopping criterion for tractography in the tumor region.

### Tractography

Tractography results are presented in Figures 5–6. Each figure shows a different subject (PAT05 and PAT26), whereas each row displays a different slice through a part of the tumor region. The slices in the first rows match those shown in Figures 1 and 3, respectively. Columns again directly compare the three different CSD pipelines’ results. Overall, outcomes reflect the quality of the respective FODs which guided the tractography as expected. ST-CSD guided tractography shows streamlines in tumor regions, but they are somewhat sparse and noisy at times due to the many false positive FOD lobes. The FOD amplitude threshold struggles (and often fails) to separate valid structure from noise within and between FODs. As shown before, the performance is similarly limited in cortical and other genuine GM areas. On the other hand, MSMT-CSD guided tractography strictly misses most WM tracts in the entire tumor region. While this happens despite lowering the FOD amplitude threshold during tractography, it is not surprising given that MSMT-CSD produces no FOD at all for a large number of tumor voxels. Finally, SS3T-CSD guided tractography successfully recovers coherent WM tracts within the tumors, which integrate well with surrounding (healthy) WM anatomy. Also note that some tumor areas in patient PAT26 (Figure 6) contain no streamlines at all. However, this is consistent with exceptionally hypo-intense regions on the T1-weighted image as well as very high CSF-like (fluid-like) content in the SS3T-CSD tissue map (Figure 3).

**Figure 5.**
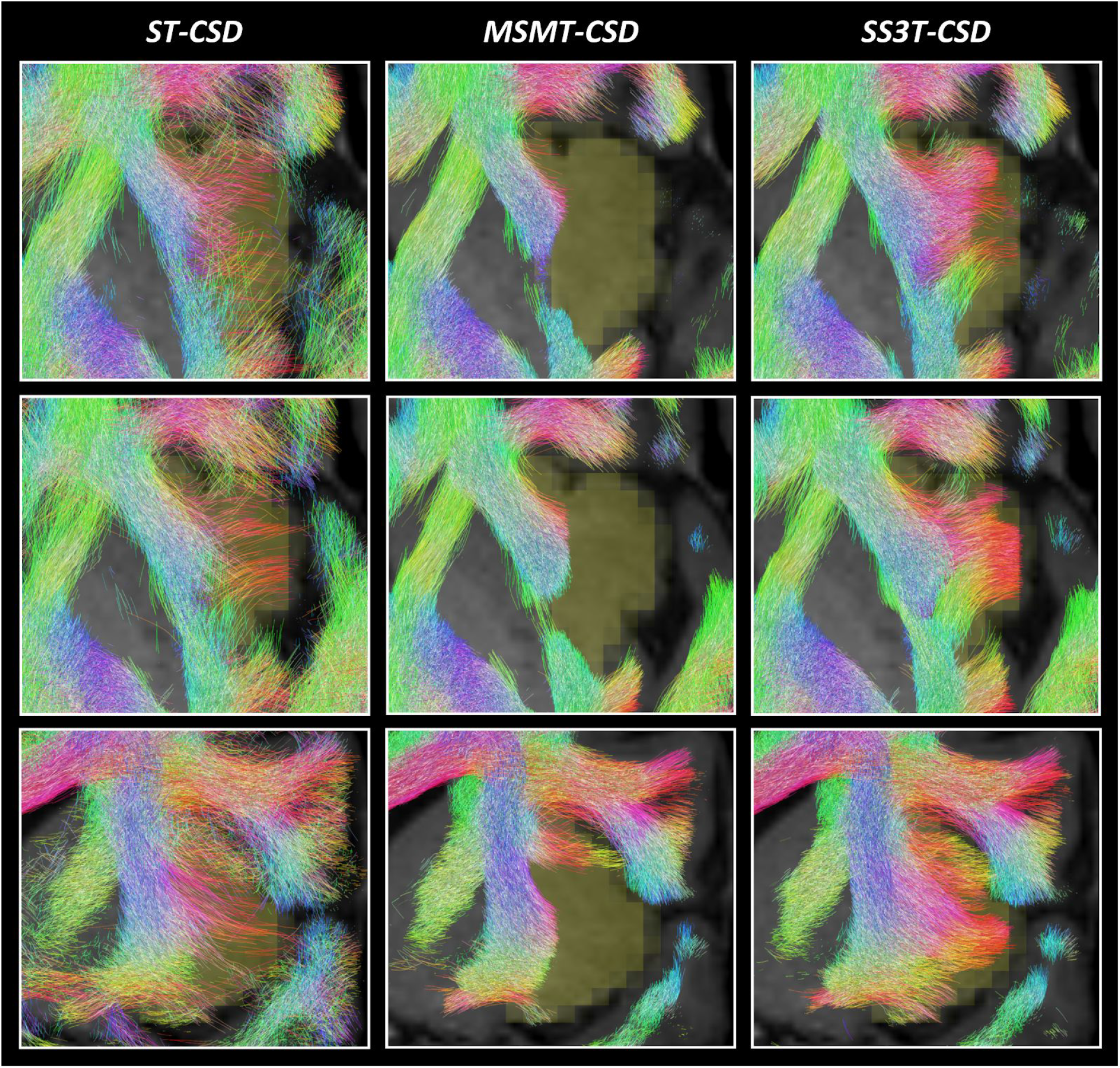
Comparison of tractography results based on ST-CSD, MSMT-CSD and SS3T-CSD pipelines for patient PAT05 (oligo-astrocytoma WHO grade II). Each result is overlaid on the T1-weighted image and the tumor segmentation is shown in yellow (at the spatial resolution of the dMRI data). Streamlines are colored using the DEC convention (red: left-right; green: anterior-posterior; blue: superior-inferior) and shown within a 2.5 mm thick “slab” centered around the slice. Each row shows a different slice. Top row: same axial slice as shown in Figure 1. Middle row: axial slice directly below the previous slice. Bottom row: coronal slice through the tumor volume.

**Figure 6.**
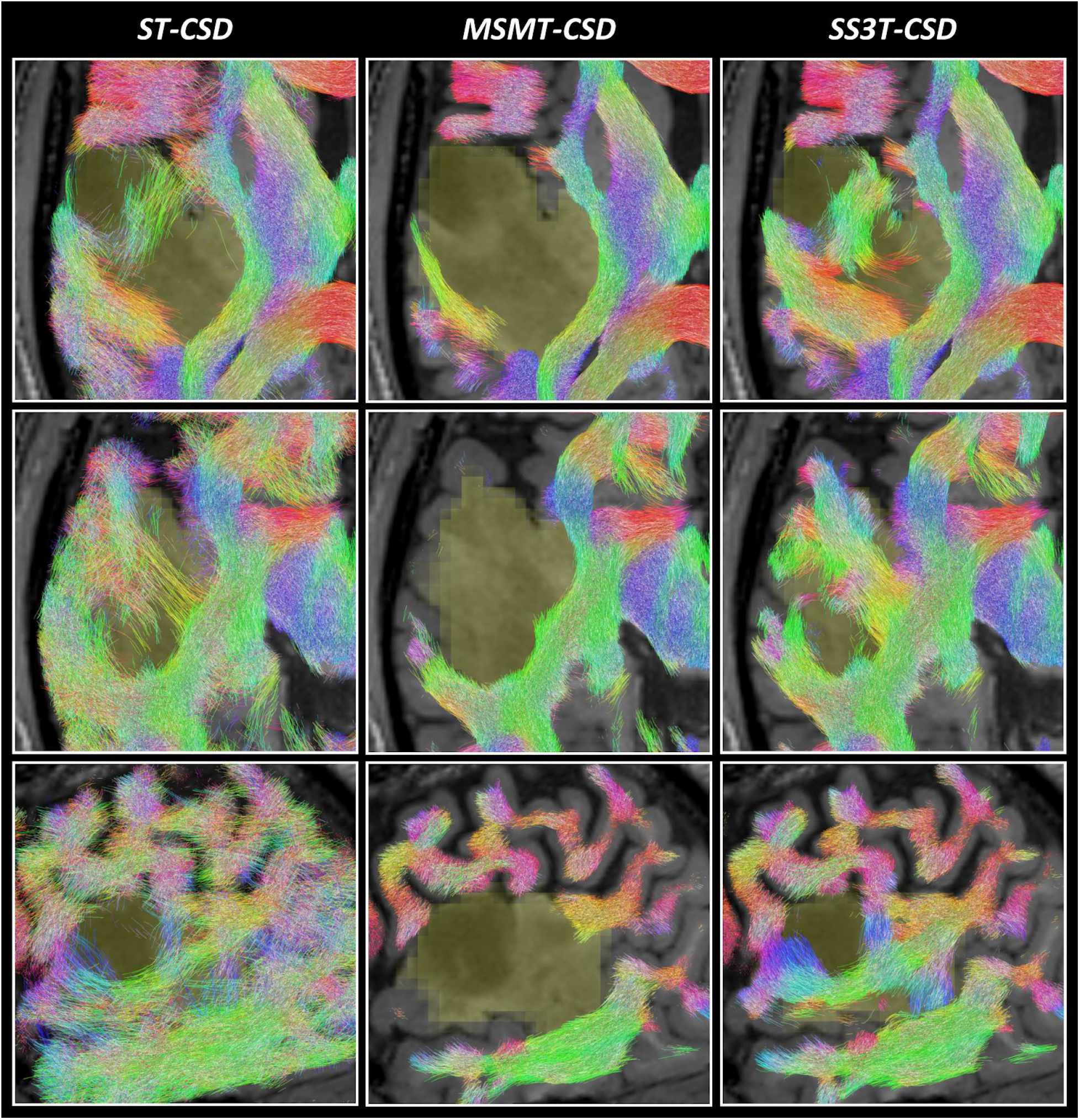
Comparison of tractography results based on ST-CSD, MSMT-CSD and SS3T-CSD pipelines for patient PAT26 (anaplastic astrocytoma WHO grade III). Each result is overlaid on the T1-weighted image and the tumor segmentation is shown in yellow (at the spatial resolution of the dMRI data). Streamlines are colored using the DEC convention (red: left-right; green: anterior-posterior; blue: superior-inferior) and shown within a 2.5 mm thick “slab” centered around the slice. Each row shows a different slice. Top row: same axial slice as shown in Figure 3. Middle row: other axial slice, further down the tumor volume. Bottom row: sagittal slice through the tumor volume.

## Discussion

In this study, we evaluated the performance of different CSD techniques for the purpose of identifying white matter tracts close to and within brain tumors. This depends on each CSD variant’s ability to resolve the distributions of white matter fiber orientations (WM FODs) in the tumor region, which are then used directly to guide tractography algorithms. We found that each CSD technique performed very differently at this challenge, each showing distinct characteristics revealing relevant strengths and limitations.

### Single-tissue CSD still has major limitations

Whereas previous studies found that single-tissue CSD substantially improved tractography results for presurgical planning compared to diffusion tensor imaging (DTI) (Farquharson et al., 2013; Küpper et al., 2015; Mormina et al., 2016), major limitations clearly still remain. Single-tissue CSD performs well in healthy WM, thus allowing improved tractography over DTI in the unaffected WM of the brain, which in turn can help to identify and locate important tracts close to the tumor. However, single-tissue CSD is severely limited in its capacity to resolve reliable WM FODs in the presence of other tissues. This has already been demonstrated before to be the case for regions of gray matter (GM), and was originally addressed by the introduction of MSMT-CSD (Jeurissen et al., 2014). Our results showed that single-tissue CSD faces a very similar limitation within infiltrating tumors. Specifically, resulting FODs might show *some* structure and slight spatial coherence, but they are severely distorted by random noisy lobes which often even completely swamp the underlying anisotropic structure of WM tracts within the tumor. These noisy lobes can be understood as isotropic diffusion signal contributions from other tissues “polluting” the WM FODs. The consequential limitations for tractography are also similar to those observed, for example, in GM: it essentially becomes impossible to define a single threshold which separates all noisy lobes from those related to genuine structure. This results in a number of both false positive streamlines as well as missing structures. This makes it hard—if not impossible—to safely determine whether (and what) parts of WM bundles exist within the tumor using single-tissue CSD guided tractography (Mormina et al., 2015). Finally, in the vicinity of healthy GM, single-tissue CSD guided tractography will of course also suffer the (same) limitations previously shown (Jeurissen et al., 2014).

### MSMT-CSD underestimates and misses within-tumor white matter tracts

In line with previous findings, MSMT-CSD improves tractography in healthy regions of GM, while maintaining the existing benefits of ST-CSD in healthy WM (Jeurissen et al., 2014). These improvements in GM regions were expected and can be explained by contributions of the GM compartment in the model, resulting in removal of most false positive WM FOD lobes. While not unique to the scenario of a brain tumor, this itself is an important benefit, as a tumor may be located close to or partially within the GM and the quality of tractography in this area might have an impact on the surgical approach. However, WM FODs were severely underestimated in large parts of the tumors we examined, up to the point they were entirely absent from most voxels in the tumor regions. Particularly in a presurgical setting, this introduces a severe risk of causing damage to functional parts of WM tracts. When assessing the general nature of the tumor, MSMT-CSD results might even appear to suggest that a genuinely infiltrative tumor is not infiltrating at all. In all cases where the WM FOD was underestimated or absent, we found a strong presence of GM-like diffusion signal contributions. That is, the b-value dependent contrast of the diffusion signal had supposedly dominated the fit, while the anisotropy in the signal was in turn partially or entirely ignored. The consequences for tractography are obvious, as most of the infiltrated WM tracts are not recovered. Lowering the FOD amplitude threshold in tractography cannot overcome this limitation, as the FODs are simply absent in a majority of voxels. These findings make it hard to “strictly” recommend MSMT-CSD over ST-CSD for the purpose of presurgical planning, especially considering the higher acquisition requirements: while ST-CSD suffers from noisy WM FODs within the tumor region, some anisotropic structure could sometimes still be observed (although not too reliably so).

### SS3T-CSD successfully recovers within-tumor white matter structure

As shown in previous work, SS3T-CSD maintains the benefits of both ST-CSD and MSMT-CSD in healthy WM tissue and improves WM FOD estimation and tractography in healthy regions of GM in a similar fashion as MSMT-CSD, yet relying only on single-shell (+b=0) dMRI data (Dhollander and Connelly, 2016). Similar to MSMT-CSD, the GM compartment in the model mostly explains and enables the removal of false positive WM FOD lobes in GM areas. However, in tumor regions, SS3T-CSD results show a striking difference compared to those obtained by MSMT-CSD. Whereas MSMT-CSD severely underestimates the presence of WM FODs or entirely removes them, WM FODs are successfully recovered by SS3T-CSD with spatially varying amounts of presence across the entire tumor volumes. Compared to the noisy within-tumor WM FODs from ST-CSD though, those obtained from SS3T-CSD reveal clear *structure* which is spatially coherent as well as consistent with known and surrounding anatomy. Furthermore, this is also consistent with the notion that gliomas show various degrees of infiltration in healthy WM structures.

As mentioned before, SS3T-CSD achieves this feat using only the single-shell (+b=0) part of the data. At first sight, this might seem paradoxical, but choosing to use this particular part of the data proves to be one of its very strengths: tumor tissue no longer appears entirely GM-like (under the assumptions and constraints of MSMT-CSD), and the anisotropy in the single diffusion weighted shell is successfully fitted by the WM FOD. However, large degrees of infiltration of tumor tissue, possibly complemented by other damage to the WM tracts, result in lower WM FOD amplitudes. This is entirely sensible, but introduces a challenge for tractography algorithms which rely on the FOD amplitude as a means to stop streamlines venturing too far or deep into non-WM areas. In this work, we addressed this challenge by gradually lowering the FOD amplitude threshold during tractography when approaching the tumor, as well as further within the tumor. For presurgical planning, this is a feasible solution, as a segmentation of the tumor will naturally be part of the process. Even so, the threshold has to be carefully managed to avoid missing genuine structure and WM tracts. Therefore, we would highly recommend complementing tractography findings with an assessment of the “underlying” WM FODs in order to avoid overlooking relevant features.

Compared to ST-CSD and MSMT-CSD, we conclude that SS3T-CSD provides strict improvements for our data. Relative to ST-CSD, WM FODs are far less noisy and generally “cleaned up” by removing diffusion signal contributions from other tissues. Relative to MSMT-CSD, within-tumor WM FODs are successfully recovered, and this while not depending on the increased acquisition requirements of MSMT-CSD. The complete multi-shell protocol took 15 minutes to acquire, but the single-shell (+b=0) subset of the data would only have required an acquisition time of about 8 minutes. With recent developments in MRI acquisition techniques such as simultaneous multi-slice imaging, this can even be further reduced to about 3 minutes, e.g. using a multiband factor of 3 (Feinberg and Setsompop, 2013).

### Limitations and future directions

A limitation of our current work lies in the fact that we only had access to data of patients with gliomas of WHO grade II and III. Nevertheless, our results carefully suggest that the general patterns observed in our findings might apply to other types of infiltrating tumors or other pathological tissue compositions. Previous works have observed similar benefits of SS3T-CSD in white matter hyperintensities (Dhollander et al., 2017; Mito et al., 2018; Mito et al., 2019), which occur in the brain with healthy aging as well as for example in Alzheimer’s disease. Future work could aim to extend these findings to other tumor types and pathologies. The aforementioned works have also started to look at the heterogeneity of tissue content within white matter hyperintensities. Accordingly, this could be investigated in different types of tumors, but interpretation of such results is certainly non-trivial and should be approached with great care. Our findings in this work suggest that a (spatially varying) combination of GM-like and CSF-like (fluid-like) diffusion signal is able to fit at least a substantial part of the single-shell (+b=0) diffusion signal resulting from infiltrating tumor tissue in our data, as evidenced by the WM FOD structure that remains once the other tissue compartments are filtered out. To further assess and eventually interpret the microstructural contents of tumors, 3-tissue CSD results could be complemented with information from other diffusion models as well as other types of diffusion acquisitions (encodings) and even other MRI modalities (Nilsson et al., 2018; Szczepankiewicz et al., 2016). Although tumor tissue characterization is a separate goal from presurgical planning, it can inform the decision on whether to perform surgery in the first place.

While we have shown that SS3T-CSD improves the reconstruction of within-tumor WM FODs, important challenges for tractography algorithms remain. We were able to address one such challenge—the overall lower within-tumor WM FOD amplitude—relatively well with a pragmatic solution, but tractography is still an ill-posed problem. One of the main problems with tractography algorithms in general, is a tendency towards a large proportion of false positive streamlines (Maier-Hein et al., 2017). It should be noted, however, that this particular problem is of slightly lesser concern for presurgical planning, as the primary focus in this context typically entails the preservation of a set of well-defined bundles. Such prior information is often actively included by means of “targeted” tractography, which may for example require that tracts traverse specific predefined regions. Furthermore, the assessment of these tractography results is also done by experts with a deep knowledge of known brain anatomy; hence this is not merely an automated “blind” approach.

False positive streamlines and tracts are however of greater concern to certain non-targeted (automated) whole-brain analysis techniques, such as typical connectomics pipelines. Different approaches are being proposed to tackle this long-standing challenge, including machine learning techniques that are pretrained with a comprehensive set of known (anatomically valid) WM tracts (Wasserthal et al., 2018) and model-driven strategies which try to explain the data using a sparse set of tracts and other priors (Schiavi et al., 2019). Improvements in the reconstruction of within-tumor WM FODs, such as those achieved by SS3T-CSD, can directly provide a more reliable starting point for these advanced tractography strategies.

Although the challenges related to tumor infiltration thus might be effectively addressed, tumor mass effects on the other hand might prove to pose unique and non-trivial challenges to pretrained machine learning strategies, such as (Wasserthal et al., 2018) in particular, as these often partially rely on an expected location, shape or size of specific WM bundles. While beyond the scope of our work, in-depth evaluation of what kinds of features certain techniques—including more complex machine learning strategies—rely upon to perform robustly is an interesting avenue for future research. Similarly in the presurgical planning setting, mass effects might effectively end up being one of the final main challenges to address. As such, any scenario that relies on prior knowledge (be it machine learned or acquired through human experience) will be challenged when extreme deformation and displacement of structures takes place. To add to the challenge, techniques for accurate compensation for brain shift during surgery in order to allow for a robust and continuous alignment of the patient to their preoperative images, are not yet established (Gerard et al., 2017). As a result, improving tractography as well as interventional alignment systems are both active fields of ongoing research.

## Conclusion

We found that 3-tissue CSD pipelines can improve greatly over the original single-tissue CSD for the purpose of FOD reconstruction and tractography within (and close to) tumor regions. However, even though relying on greater acquisition requirements, MSMT-CSD introduced a distinct risk to *underestimate* the presence of intact WM tracts within infiltrating tumors. Perhaps surprisingly, SS3T-CSD was able to provide a far more complete reconstruction, even though relying on less data. This provides a unique opportunity to improve clinical practice in the context of presurgical planning with minimal requirements.

## Acknowledgements

This project has received funding from the Special Research Funds (BOF) of the University of Ghent (01MR0210 and 01J10715), Grant P7/11 from the Interuniversity Attraction Poles Program of the Belgian Federal Government, and the European Union’s Horizon 2020 Framework Programme for Research and Innovation under the Specific Grant Agreement No. 785907 (Human Brain Project SGA2).

We are grateful to the National Health and Medical Research Council (NHMRC) of Australia and the Victorian Government’s Operational Infrastructure Support Program for their support.

We would like to thank Prof. Dr. Dirk Van Roost, Prof. Dr. Eric Achten, Stephanie Bogaert, Robby De Pauw, Hannes Almgren, Iris Coppieters, Jeroen Kregel, Mireille Augustijn and Helena Verhelst for their help in acquiring the data.

## Footnotes

1 The term “tissue” is used in an abstract sense in context of these multi-tissue CSD techniques: it refers in general to extra signal compartments in the model, even e.g. (cerebrospinal) fluid.

